# Tracking-based rolling angles recovery method for holographic tomography of flowing cells

**DOI:** 10.1101/2021.06.01.446558

**Authors:** Daniele Pirone, Pasquale Memmolo, Francesco Merola, Lisa Miccio, Martina Mugnano, Amedeo Capozzoli, Claudio Curcio, Angelo Liseno, Pietro Ferraro

**Affiliations:** Institute of Applied Sciences and Intelligent Systems (ISASI), National Research Council (CNR) of Italy, Via Campi Flegrei 34, 80078 Pozzuoli, NA, Italy; Department of Electrical Engineering and Information Technology, University of Naples “Federico II”, Via Claudio 21, 80125 Naples, Italy

**Keywords:** Digital Holography, Holographic Tomography, Tomographic Flow Cytometry, 3D Holographic Particle Tracking, Rolling Angles Recovery, Phase Image Similarity Metric, Cell Phantom, Tumor Cell

## Abstract

Holographic Tomography (HT) is an emerging label-free technique for microscopic bioimaging applications, that allows reconstructing the three-dimensional (3D) refractive index (RI) distribution of biological specimens. Recently, an in-flow HT technique has been proposed in which multiple digital holograms are recorded at different viewing angles around the sample while it flows and rotates within a microfluidic channel. However, unlike conventional HT methods, there is no a priori information about cell 3D orientations, that are instead requested to perform any tomographic algorithm. Here we investigate a tracking-based rolling angles recovery method, showing robustness against the sample’s features. It is based on a phase images similarity metric recently demonstrated, that exploits the local contrast phase measurements to recognize a full cell rotation within the microfluidic channel. Hence, the orientations of the flowing cells are retrieved from their positions, which are in turn computed through the 3D holographic tracking. The performances of the rolling angles recovery method have been assessed both numerically, by simulating a 3D cell phantom, and experimentally, by reconstructing the 3D RI tomograms of two cancer cells. Both the numerical and the experimental analysis have been performed at different spatial resolutions. This rolling angles recovery method, not depending on the cell shapes, the RI contents, and the optical experimental conditions, could pave the way to the study of circulating tumor cells (CTCs) in the challenging tool of liquid biopsy.

## 1. Introduction

Biological samples are transparent objects that mainly perturb the phase component rather than the amplitude one of an incident optical radiation^1^. Fluorescence Microscopy (FM) has established as the main imaging tool in cell biology since stains or fluorescent tags are used to make them visible within amplitude images^1,2^. However, FM is often qualitative and limited by photobleaching, i.e. the reduction of the fluorescence intensity due to the photochemical degradation of the fluorophore, and by the phototoxicity of the fluorescent proteins or dyes, which can alter the normal cell physiology^1^. These drawbacks have been overcome by Digital Holography (DH), which allows recording both the amplitude and phase information into interference fringe patterns in two-dimensional (2D) images^3^. Then, from the digital hologram, a Quantitative Phase Map (QPM) can be recovered through numerical phase retrieval techniques^3^. A QPM can be physically interpreted as the integral image of the sample’s RI contrast along the optical *z*-axis. To decouple the spatial RI distribution from the sample’s *z*-thickness, multiple QPMs must be recorded at different viewing angles around the sample. This is the principle behind HT, which is a label-free optical microscopy technique that allows reconstructing the 3D RI distribution of a biological specimen^4^. Therefore, HT is finding various applications in biomedicine thanks to its unique property of providing a 3D quantitative analysis at the single-cell level about morphologies and RI statistics, without limitations due to the use of dyes or markers^1,4^. To perform HT, two conventional methods can be exploited. In the first one, the fixed sample is scanned by the incident optical field at different beam directions^5-8^. In the second one, the sample is rotated under a fixed beam direction through mechanical^9^ or optical^10^ forces. However, in the first case the maximum illumination angle typically cannot exceed 150°, thus limiting the tomographic accuracy, while in the second case the sample can be perturbed by the external forces needed to rotate it, and the complex recording system cannot guarantee a high throughput analysis^11^. Instead, an alternate HT technique has been proposed, i.e. tomographic flow cytometry by DH^12^, in which a DH microscope records, along a fixed direction, the digital holograms of cells while flowing within a microfluidic channel and rotating due to the hydrodynamic forces produced by a laminar flow, thus providing full sample rotation, no external alteration, and potentially high-throughput analysis of single cells. However, unlike the conventional HT techniques, the 3D positions and the viewing/rolling angles are not a priori known, thus they must be retrieved to implement a tomographic algorithm. Cell 3D positions can be computed through assessed 3D holographic particle tracking algorithms^13^. Instead, the existing solutions to estimate the unknown rolling angles of flowing cells in microfluidic environment were tailored to the shape and RI content of the observed sample^12,14^. However, generally this information is unavailable, because it is the outcome of the HT analysis. Recently, we proposed a tracking-based rolling angles recovery method, and we demonstrated its robustness against the sample’s features^15^. A full cell rotation is identified by an ad hoc phase image similarity metric, from which all the other unknown rolling angles are retrieved thanks to the microfluidic properties of the microchannel. Here we investigate the robustness of this method against the reconstruction pixel dimension, in order to assess its feasibility in several DH experimental conditions. In fact, in some applications, the spatial resolution must be sacrificed to the advantage of a greater field of view (FOV). To this aim, we simulate a 3D flowing and rotating cell phantom, used as ground truth, and we reconstruct the 3D RI tomograms of two CTCs, belonging to the CHP-134 neuroblastoma cell line, recorded with different microscope objectives.

## 2. Materials and Methods

The HT recording system, sketched in Fig. 1(a), is based on a DH microscope, that exploits a Mach-Zehnder interferometer in off-axis configuration. The light beam generated by a fiber couple laser (532 nm – 400 mW) is divided by a beam splitter into an object beam and a reference beam. The object beam illuminates the biological sample, which mainly changes its phase component, and, after passing through a microscope objective (oil-immersion, Plan-Apochromat), it interferes with the reference beam. The resulting interference fringe pattern propagates up to a CMOS camera (USB 3.0 u eye from IDS - 2048×2048 – pixel size 5.5 μm), which records the digital holograms at 35 fps. The biological sample is illuminated while flowing and rotating within a microfluidic channel (PMMA polymeric material– length 7 cm – cross section 200 μm × 200 μm), thanks to the parabolic velocity profile of the laminar flow^16^ created by a microfluidic pump (CETONI – neMESYS) pushing at 7 nL/s. The microfluidic properties ensure that cells flow along the *y*-axis and continuously rotate around the *x*-axis, according to the reference system reported in Fig. 1(a). Each hologram of the recorded sequence is demodulated by extracting the real diffraction order through a band-pass filter, because of the off-axis configuration^3^. Then, a holographic tracking algorithm^13^ is used to estimate the 3D positions of the flowing cells along the microfluidic channel. It consists of two successive steps. The first one is the axial *z*-localization, in which the hologram is numerically propagated at different *z*-positions through the Angular Spectrum formula^17^, and, for each of them, the Tamura Coefficient^13^ (TC) is computed on the region of interest (ROI) containing the cell within the amplitude of the reconstructed complex wavefront. By minimizing this contrast-based metric, the cell *z-*position in each frame can be recovered, and the cell can be refocused. After computing the in-focus complex wavefront, the corresponding QPM is obtained by performing the phase unwrapping algorithm^18^. The second holographic tracking’s step is the transversal *xy*-localization, which is obtained by computing the weighted centroid of the cell in its QPM^13^. The 3D holographic tracking allows centering each cell in all the QPM-ROIs of their recorded rolling sequence, thus avoiding motion artefacts in the successive tomographic reconstruction. Moreover, the *y*-positions can be exploited to estimate the unknown rolling angles, according to the strategy proposed in Ref. 15. A phase image similarity metric, namely Tamura Similarity Index (TSI), based on the evaluation of the local contrast by TC, is computed on all the QPMs of the rolling cell. It has been demonstrated minimizing in the frame *f*_*180*_ at which a 180° of rotation with respect to the first frame of the sequence has occurred. Thanks to the above-mentioned microfluidic properties and to the high recording frame rate, a linearization of the relationship between the angular and the translational speeds can be assumed. Hence, all the *N* unknown rolling angles are computed as follows

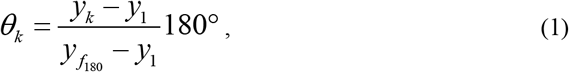

where *k=1,…,N* is the frame index. Finally, the tomographic reconstruction is performed by the Filtered Back Projection (FBP) algorithm^19^.

**Fig. 1.**
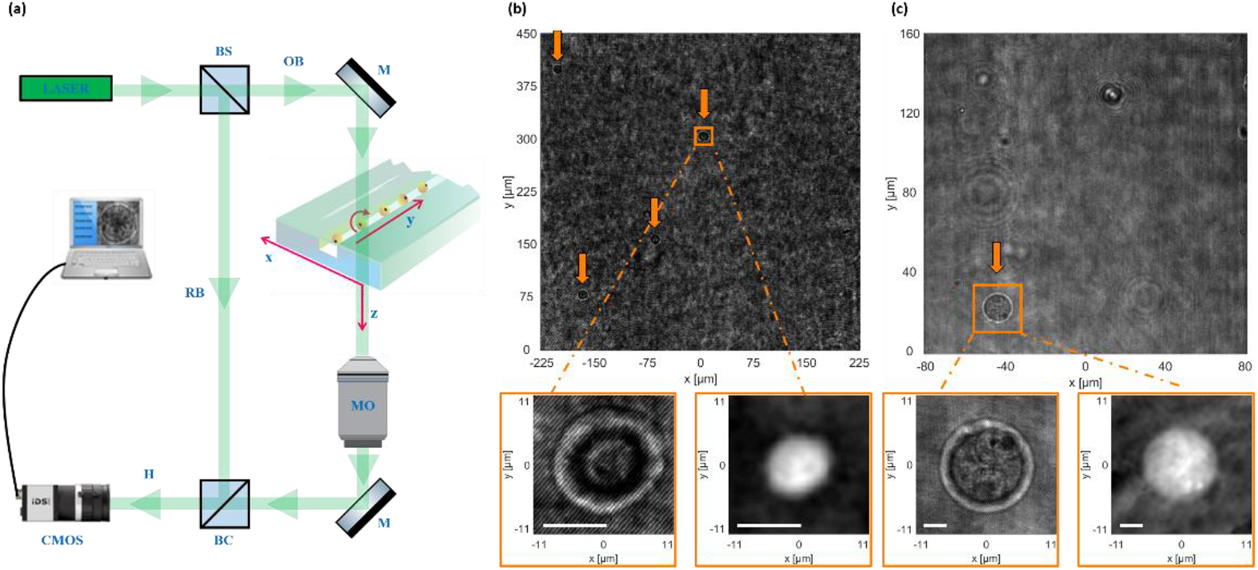
HT recording system. (a) Sketch of the DH microscope. OB, Object Beam; RB, Reference Beam; H, Hologram; MO, Microscope Objective; M, Mirror; BS, Beam-Splitter; BC, Beam-Combiner; CMOS, Camera. (b,c) Digital holograms of CHP-134 cells recorded by 20× and 40× microscope objectives, respectively. Orange arrows indicate the flowing and rotating cells. In the orange inserts, zoom of the holographic ROI (left) and corresponding QPM (right). Scale bars are 50 pixels.

In order to evaluate the trade-off between the spatial resolution and the FOV, CHP-134 cells have been recorded by using two different microscope objectives with 20× and 40× nominal magnifications, as shown in Figs. 1(b,c), respectively. Standard USAF targets are used to measure the effective magnifications and consequently the reconstruction pixel dimensions in the two experimental conditions, which are 0.22 μm for the nominal 20× and 0.08 μm for the nominal 40×. Obviously, the 40× microscope objective allows analysing a smaller FOV with a better spatial resolution (FOV_40×_ = 0.03 mm^2^), as highlighted by the holographic ROIs and the corresponding QPMs in the inserts in Figs. 1(b,c). However, in some applications, to make a statistical analysis on a large number of cells, a greater FOV is preferred. In particular, the 20× microscope objective is particularly suitable for high-throughput purposes (FOV_20×_ = 0.20 mm^2^) but to the detriment of the tomographic accuracy. In fact, the tomographic accuracy deteriorates not only because of the decrease of the reconstruction pixel dimension, but also because the lower spatial resolution complicates the search of the frame *f*_*180*_, since it is based on the identification of image similarities. In order to investigate the dependence of the *f*_*180*_ accuracy to the spatial resolution, we simulate a 3D numerical cell phantom with different reconstruction pixel dimensions. The 3D numerical model contains the main four sub-cellular structures, i.e. the cell membrane (RI = 1.350), the cytoplasm (RI = 1.365), the nucleus (RI = 1.380), and some mitochondria (RI = 1.410), which have been simulated according to morphological and RI information reported in literature^20,21^. To obtain synthetic QPMs, the simulated 3D RI distribution is numerically integrated at equally spaced angles around the same axis in the [0-360]° interval. To evaluate the robustness of the rolling angles recovery method, we simulate 90 QPM sequences by selecting the following parameters

- 2 possible cell external shapes, i.e. spherical and non-spherical;
- 5 possible cell rolling angle steps, i.e. {3, 3.5, 4, 4.5, 5}°;
- 3 possible Gaussian distributions of the sub-cellular RIs, with standard deviations σ_RI_ = {0.01, 0.03, 0.05} and with the average values reported above;
- 3 possible Gaussian noises added to each QPM, with standard deviations σ_N_ = {0.1, 0.4, 0.7} rad and with zero-mean.

Moreover, the same 90 QPM sequences have been simulated with five different reconstruction pixel dimensions, i.e. {0.05, 0.10, 0.15, 0.20, 0.25} μm, in order to include the experimental spatial resolutions. Finally, to each QPM, a smoothing operation has been applied to simulate the point spread function of an imaging system. In Figure 2(a), a synthetic QPM simulated with non-spherical shape, σ_RI_ = 0.03, and σ_N_ = 0.4 rad is reported. In the blue box zoomed in Figs. 2(b,c), two different reconstruction pixel dimensions have been simulated, i.e. 0.05 μm and 0.25 μm, respectively. For each synthetic QPM sequence, the TSI method has been applied to find the *f*_*180*_ frame. Moreover, another state-of-the-art image similarity metric has been tested, i.e. the multi-scale structural similarity index (MS-SSIM)^22^. For each reconstruction pixel dimension, the accuracy of each metric has been computed as the percentage of correct *f*_*180*_ identifications among the 90 QPM sequences. As shown in Figure 2(d), the MS-SSIM accuracy decreases a lot with the pixel dimension, while the TSI accuracy keeps about constant above the 90 % of accuracy at different spatial resolutions. Identifying the right *f*_*180*_ frame is the key point to make a right estimation of the angular sequence, which in turn allows reconstructing the 3D RI tomogram with high accuracy. In Figure 2(e), the root mean square error (RMSE) between the estimated angular sequence and the one simulated with three different angular steps (3°, 4°, and 5°), is reported versus the *f*_*180*_ error, i.e. the difference between the *f*_*180*_ frame used in Eq. (1) to retrieve the unknown rolling angles and the true *f*_*180*_ frame. It is clear that also a ±1 frame error causes a great deviation from the true angular sequence, and this effect increases with the angular step.

**Fig. 2.**
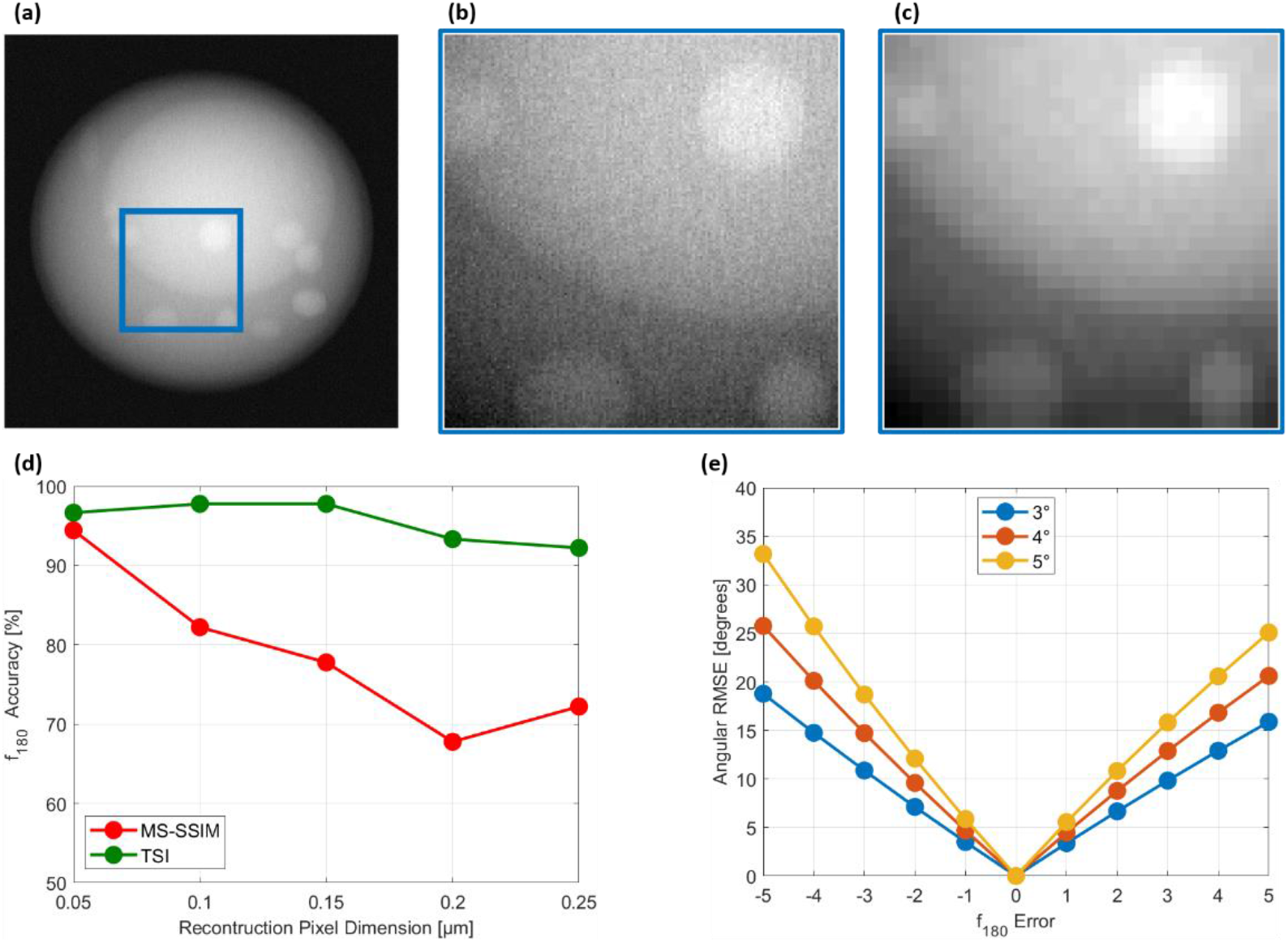
Assessment of the *f*_*180*_ search at different spatial resolutions. (a) Synthetic QPM, simulated with non-spherical shape, σ_RI_ = 0.03, and σ_N_ = 0.4 rad. The side of the blue box is 7.5 μm. (b,c) Zoom of the blue box in (a), simulated with 0.05 μm and 0.25 μm reconstruction pixel dimensions, respectively. (d) Percentage of correct *f*_*180*_ frames identified by TSI (green) and MS-SSIM (red) at different reconstruction pixel dimensions. (e) RMSE between the angular sequences simulated with angular steps of 3° (blue), 4° (orange), and 5° (yellow), and the angular sequences estimated with different *f*_*180*_ frames. The *f*_*180*_ error is the difference between the *f*_*180*_ frame used in Eq. (1) to retrieve the unknown rolling angles and the true *f*_*180*_ frame.

## 3. Results and Conclusions

The rolling angles recovery method has been employed to estimate the unknown viewing angles of the two CHP-134 cells highlighted in Figs. 1(b,c). To this aim, for each frame of the QPM sequence, the TSI has been computed, which minimizes at frame *f*_*180*_. As expected, the TSI trend appears more regular in the 40× case in Fig. 3(d) than the 20× case in Fig. 3(a), because of the higher spatial resolution in the corresponding QPMs. After using the computed *f*_*180*_ value in Eq. (1), the FBP algorithm has been implemented to reconstruct the 3D RI tomograms. In Figures 3(e,f), respectively the isolevels representation and the central slice of the CHP-134 cell recorded by the 40× microscope objective show a much more detailed inner structure with respect to the CHP-134 cell recorded by the 20× microscope objective, displayed in Figs. 3(b,c). However, as demonstrated by the numerical simulation, the TSI allows a great robustness against the loss of resolution due to the employment of microscope objectives with low nominal magnifications. This property is fundamental to access the 3D RI information of flowing cells regardless the experimental conditions. In this way, the tomographic flow cytometry becomes a powerful tool to be exploited for different purposes. For example, with a high spatial resolution, a detailed cell characterization could be performed. At the same time, a lower spatial resolution is more suitable for high-throughput analysis. Recently, a classification task has been solved to discriminate different CTC lines by using the 2D holograms of flowing cells^23^. The next challenge is the development of machine-learning tools fed by a large number of 3D RI tomograms, from which more informative features can be extracted with respect to the 2D imaging, thus covering a further step towards the implementation of the liquid biopsy paradigm^24^.

**Fig. 3.**
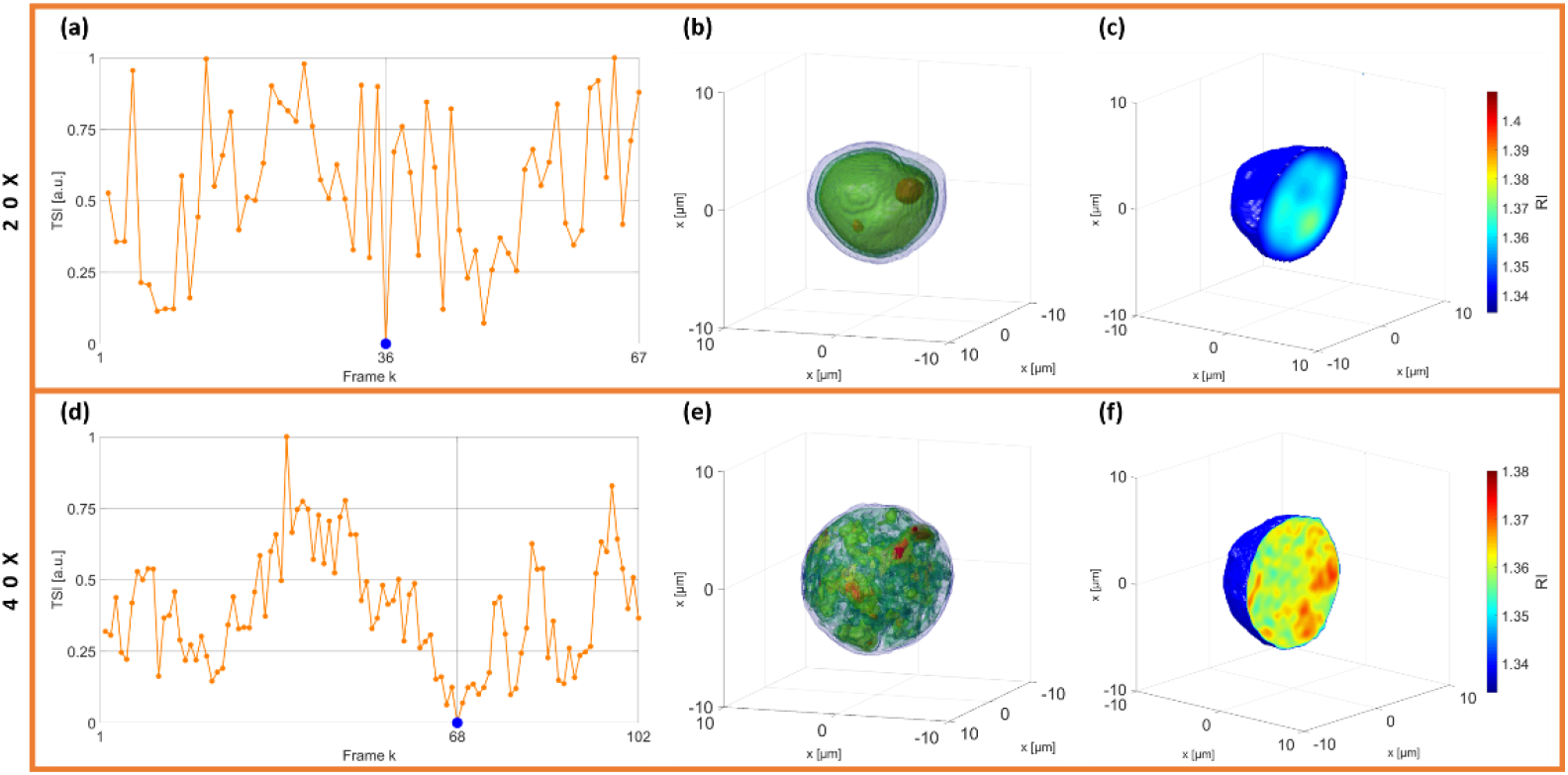
HT reconstructions of two CHP-134 cells recorded by 20× (a-c) and 40× (d-f) microscope objectives. (a,d) TSI computed for each QPM (orange dots) to find the *f*_*180*_ frame (blue dot). (b,e) Isolevels representation and (c,f) central slice of the 3D RI tomogram reconstructed by in-flow HT. In (b,c), each side of the 3D array is 90 pixels. In (c,f), each side of the 3D array is 250 pixels.

## References

1. Park, Y., Depeursinge, C. and Popescu, G., “Quantitative phase imaging in biomedicine,” Nat. Photonics 12(10), 578–589 (2018).

2. Diaspro, A., “Optical Fluorescence Microscopy,” Springer (2011).

3. Kim, M. K., “Principles and techniques of digital holographic microscopy,” SPIE Rev. 1, 018005–018048 (2010).

4. Jin, D., Zhou, R., Yaqoob, Z. and So, P., “Tomographic phase microscopy: Principles and applications in bioimaging,” J. Opt. Soc. Am. B 34(5), B64–B77 (2017).

5. Choi, W., Fang-Yen, C., Badizadegan, K., Oh, S., Lue, N., Dasari, R. R. and Feld, M. S., “Tomographic phase microscopy,” Nat. Methods 4, 717–719 (2007).

6. Kim, K., Kim, K. S., Park, H., Ye, J. C. and Park, Y., “Real-time visualization of 3-D dynamic microscopic objects using optical diffraction tomography,” Opt. Express 21, 32269–32278 (2013).

7. Isikman, S. O., Bishara, W., Mavandadi, S., Yu, F. W., Feng, S., Lau, R. and Ozcan, A., “Lens-free optical tomographic microscope with a large imaging volume on a chip,” Proc. Natl. Acad. Sci. U.S.A. 108, 7296–7301 (2011).

8. Pégard, N. C., Toth, M. L., Driscoll, M. and Fleischer, J. W. “Flow scanning optical tomography,” Lab Chip 14, 4447–4450 (2014).

9. Charrière, F., Marian, A., Montfort, F., Kuehn, J. and Colomb, T., “Cell refractive index tomography by digital holographic microscopy,” Opt. Lett. 31, 178–180 (2006).

10. Habaza, M., Gilboa, B., Roichman, Y. and Shaked, N. T. “Tomographic phase microscopy with 180° rotation of live cells in suspension by holographic optical tweezers,” Opt. Lett. 40, 1881–1884 (2015).

11. Kuś, A., Krauze, W., Makowski, P. L. and Kujawińska, M., “Holographic tomography: hardware and software solutions for 3D quantitative biomedical imaging,” ETRI Journal 41(1), 61–72 (2019).

12. Merola, F., Memmolo, P., Miccio, L., Savoia, R., Mugnano, M., Fontana, A., D’Ippolito, G., Sardo, A., Iolascon, A., Gambale, A. and Ferraro, P., “Tomographic flow cytometry by digital holography,” Light Sci. Appl. 6, e16241 (2017).

13. Memmolo, P., Miccio, L., Paturzo, M., Caprio, G. D., Coppola, G., Netti, P. A. and Ferraro, P., “Recent advances in holographic 3d particle tracking,” Adv. Opt. Photon. 7, 713–755 (2015).

14. Villone, M. M., Memmolo, P., Merola, F., Mugnano, M., Miccio, L., Maffettone, P. L. and Ferraro, P., “Full-angle tomographic phase microscopy of flowing quasi-spherical cells,” Lab Chip 18(1), 126–131 (2018).

15. Pirone, D., Memmolo, P., Merola, F., Miccio, L., Mugnano, M., Capozzoli, A., Curcio, C., Liseno, A. and Ferraro, P. “Rolling angles recovery of flowing cells in holographic tomography exploiting the phase similarity,” Appl. Opt. 60(4), A277–A284 (2021).

16. Torino, S., Iodice, M., Rendina, I., Coppola, G. and E. Schonbrun, “A microfluidic approach for inducing cell rotation by means of hydrodynamic forces,” Sensors 16(8), 1326 (2016).

17. Schnars, U. and Jüptner, W. “Digital recording and numerical reconstruction of holograms,” Meas. Sci. Technol. 13(9), R85–R101 (2002).

18. Bioucas-Dias, J. M. and Valadao, G. “Phase Unwrapping via graph cuts,” IEEE Trans. Image Process.16(3), 698–709 (2007).

19. Kak, A.C.; Slaney, M. “Principles of Computerized Tomographic Imaging,” IEEE Press, New York, NY, 49–75 (1988).

20. Wen, Y., Chen, Z., Lu, J., Ables, E., Scemama, J.-L., Yang, L., Lu, J. and Hu, X. H. “Quantitative analysis and comparison of 3D morphology between viable and apoptotic MCF-7 breast cancer cells and characterization of nuclear fragmentation,” PLoS ONE 12(9), e0184726 (2017).

21. Haseda, K., Kanematsu, K., Noguchi, K., Saito, H., Umeda, N. and Y. Ohta, “Significant correlation between refractive index and activity of mitochondria: single mitochondrion study,” Biomed. Opt. Express 6, 859–869 (2015).

22. Wang, Z., Simoncelli, E. P., Bovik, A. C. “Multiscale Structural Similarity for Image Quality Assessment,” The Thirty-Seventh Asilomar Conference on Signals, Systems & Computers, 1398–1402 (2003).

23. Dellipriscoli, M., Memmolo, P., Ciaparrone, G., Bianco, V., Merola, F., Miccio, L., Bardozzo, F., Pirone, D., Mugnano, M., Cimmino, F., Capasso, M., Iolascon, A., Ferraro, P. and Tagliaferri, R. “Neuroblastoma cells classification through learning approaches by direct analysis of digital holograms,” IEEE J Sel Top Quantum Electron, doi:10.1109/JSTQE.2021.3059532 (2021).

24. Miccio, L., Cimmino, F., Kurelac, I., Villone, M. M., Bianco, V., Memmolo, P., Merola, F., Mugnano, M., Capasso, M., Iolascon, A., Maffettone, P. L. and Ferraro, P. “Perspectives on liquid biopsy for label-free detection of circulating tumor cells through intelligent lab-on-chips,” View 1(3), 20200034 (2020).

